# Transformation of *Gardnerella vaginalis* with a *Bifidobacterium-Escherichia coli* shuttle vector plasmid

**DOI:** 10.1101/2025.02.14.638355

**Authors:** B. M. D. N. Kularatne, Janet E. Hill

**Affiliations:** Department of Veterinary Microbiology, University of Saskatchewan, Saskatoon, SK, Canada

## Abstract

*Gardnerella* spp. significantly influence female reproductive health, and are indicators of bacterial vaginosis, a common gynecological disorder. Lack of genetic tools for *Gardnerella* spp. is a hindrance in fully understanding their role in the vaginal microbiome and no naturally occurring plasmids have yet been identified in these organisms. The aim of this study was to transform *Gardnerella vaginalis* and characterize transformants carrying *Bifidobacterium-E. coli* shuttle vector pKO403-*lacZ*′-Sp. *G. vaginalis* ATCC 49145 was selected for protocol development based on its high growth rate, lack of restriction activity and susceptibility to spectinomycin. Low efficiency (∼10^2^ cfu/µg of plasmid DNA) but reproducible transformation was achieved. The expression of the spectinomycin resistance gene and the β-galactosidase gene of pKO403-*lacZ*′-Sp in *G. vaginalis* ATCC 49145 resulted in an increase in spectinomycin tolerance from 2 µg/ml (MIC) to >512 µg/ml, and an appreciable increase in β-galactosidase activity compared to the wild type. Plasmid copy number was determined to be ∼3 per genome copy. Plasmid was lost rapidly in the absence of spectinomycin selection, with only ∼13% of colony forming units retaining the resistant phenotype after 24 h of growth without selection. These results demonstrate that *G. vaginalis* can be transformed by electroporation and that pKO403-*lacZ*′-Sp can be maintained and its genes expressed in this host, offering a starting point for the development of genetic tools for mechanistic studies of this important member of the vaginal microbiome.

**Importance:** The healthy human vaginal microbiome is mainly dominated by *Lactobacillus* spp. An imbalance or shift in this population can lead to a gynecological disorder known as bacterial vaginosis (BV). In BV, there is a reduction in *Lactobacillus* spp. and overgrowth of mixed anaerobes and facultative bacteria including *Gardnerella* spp. The reason for this increase in the *Gardnerella* population and associated changes in the vaginal microbiota composition is yet not understood, and a lack of genetic tools is one of the major barriers to performing mechanistic research to study the biology of these clinically significant organisms. A first step in developing genetic tools is introducing foreign DNA. In this study, we have developed a protocol for transformation and identified a plasmid that can be maintained in *G. vaginalis*.

## Introduction

*Gardnerella* spp. are Gram-positive facultative anaerobes that colonize the human vaginal microbiome where they are associated with a common gynecological disorder known as bacterial vaginosis (BV) (1–4). BV is characterized by a depletion of normally dominant *Lactobacillus* spp. and an overgrowth of a mixed population of anaerobes and facultative anaerobes, including *Gardnerella* spp. Six species of *Gardnerella* have been defined, among which *G. vaginalis* (the type species for the genus) and *G. swidsinskii* are the most prevalent and abundant (5–7). *Gardnerella* spp. can be found in women with or without BV, but in the case of BV, a more diverse and abundant population of *Gardnerella* spp. is found (8). The reasons for the increase in the *Gardnerella* population and associated changes in vaginal microbiota community composition that characterize the BV dysbiosis are not yet understood. Over the last decade, whole genome sequence data has played an important role in predicting traits of *Gardnerella* spp., but this data does not provide insight into specific functions of individual genes. Despite a growing repertoire of *Gardnerella* sequence data, there is limited knowledge of the biology and the ecology of this organism, and characterization of gene products has been limited to in vitro studies of individual proteins (9–11). One major obstacle to addressing this knowledge gap is a lack of genetic tools and the consequent inability to manipulate genes and perform mechanistic studies.

The use of any genetic tool, including non-targeted approaches such as transposon mutagenesis and targeted methods for gene deletion, allelic replacement or introduction of marker genes, requires the introduction of exogenous DNA into host bacteria. The primary barriers that protect any bacterium from foreign DNA and make it genetically intractable include the physical barrier of the cell envelope, and restriction modification systems which are widespread defence mechanisms of bacteria (12). Physical barriers can be overcome by inducing competence with chemical treatment and electroporation (13–17). Electroporation serves as an effective method for transformation, even in cells characterized by thick cell walls (18). A brief exposure to high voltage renders the cell wall permeable, thereby facilitating the entry of DNA into the cell (19) after which the resulting transformants can be selected following a period of recovery (20). Strategies for avoiding restriction modification systems include selection of host strains lacking these systems or engineering the DNA to be introduced by removing restriction enzyme recognition sequences or altering methylation patterns to mimic the host’s DNA (21).

Naturally occurring plasmids are often a starting point for the development of vehicles for introduction of genetic material into bacteria (22). No plasmids have been identified to date in *Gardnerella* spp. (23), however, plasmids from other members of the family Bifidobacteriaceae have been used successfully to develop powerful genetic tools for *Bifidobacterium* spp. (24–28). One such family of plasmids are those based on *Bifidobacterium*– *Escherichia coli* shuttle vector pKKT427 (27, 28), including pKO403-*lacZ*′-Sp which contains two origins of replication, (ColE1 from *E. coli* and pTB6 from *Bifidobacterium*), a selectable marker (spectinomycin resistance gene), and a β-galactosidase gene for blue-white colony selection. The repB region from pTB6 is known to be temperature sensitive at 42 °C in *Bifidobacterium longum* (28); a feature that can be exploited in plasmid curing.

As there are currently no published methods to introduce foreign DNA into *Gardnerella*, developing a protocol for transformation and the identification of plasmids that can be maintained in this host is a priority. Therefore, the objectives of the current study were to establish a protocol for the transformation of *Gardnerella vaginalis* by electroporation, and to characterize transformants carrying plasmid pKO403-*lacZ*′-Sp in terms of plasmid copy number, spectinomycin tolerance, β-galactosidase activity and plasmid stability.

## Results

### Selection of strain for transformation

Growth characteristics of individual isolates of *Gardnerella* spp. are known to vary markedly (29). To facilitate development of a transformation protocol, we wanted to identify a strain that grew to high density relatively rapidly, and that could be grown in a CO_2_ incubator rather than requiring anaerobic conditions. Comparison of growth curves of five strains (*G. vaginalis* ATCC 49145, *G. vaginalis* ATCC 14018, *G. swidsinskii* LP 23b, *G. piotii* LP 13C and *G. leopoldii* LP 57) showed that the laboratory-adapted *G. vaginalis* ATCC 49145 and *G. vaginalis* ATCC 14018 had higher growth rates and reached an OD_600_ of 1.0 by 24 h while the three clinical isolates *G. leopoldii, G. piotii* and *G. swidsinskii* did not exceed an OD_600_ 0.5 even after 30 h in 7 % CO_2_, at 37 °C (Figure 1A).

**Figure 1.**
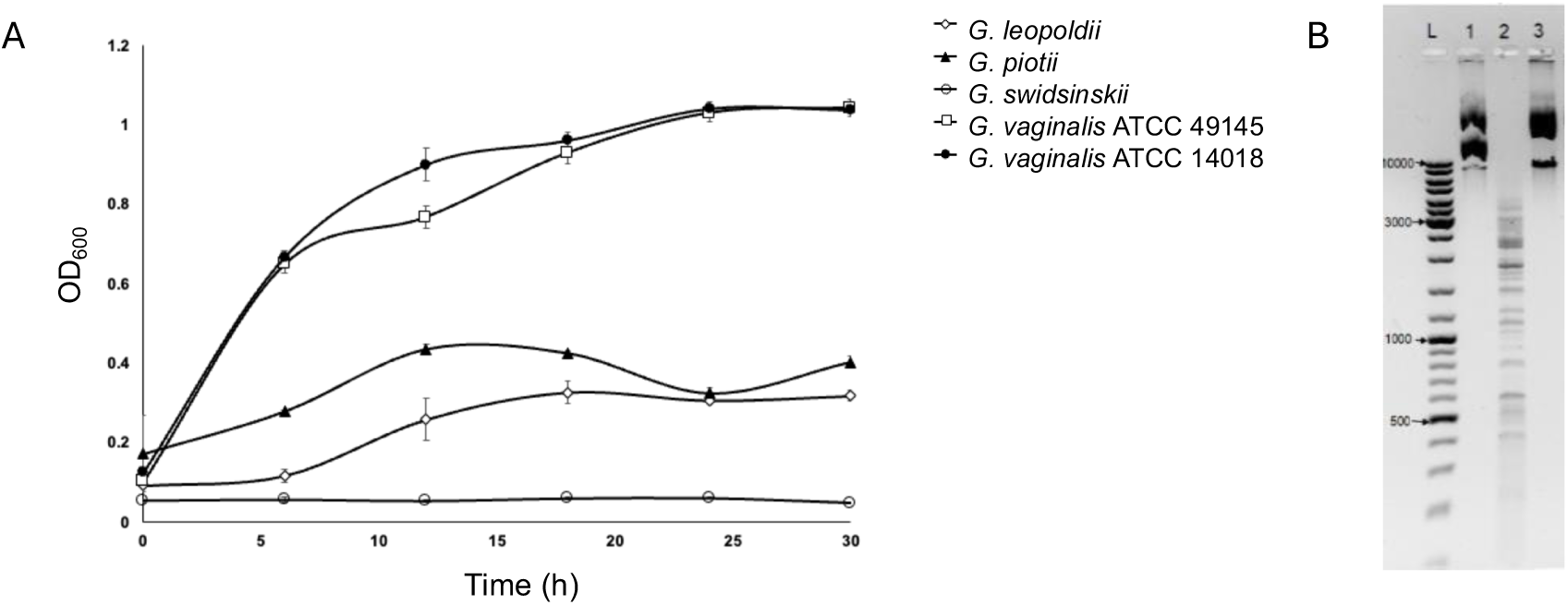
A) Growth curves of *Gardnerella* isolates at 37 °C in 7 % CO2 for 30 h. Each point represents an average of three replicates and error bars indicate standard deviation. B) Restriction activity assay results. ‘L’, Size markers; Lane 1, undigested pKO403-*lacZ′-*Sp; Lane 2, pKO403-*lacZ′-*Sp treated with the supernatant from *G. vaginalis* ATCC 14018; Lane 3, pKO403-*lacZ′-*Sp treated with the supernatant from *G. vaginalis* ATCC 49145.

In addition to robust growth, lack of restriction activity was another desirable characteristic of an isolate for use in transformation protocol development. To assess restriction activity, we incubated purified pKO403-*lacZ′-*Sp plasmid DNA with filter-sterilized culture supernatants of *G. vaginalis* ATCC 14018 and ATCC 49145. Supernatant from *G. vaginalis* ATCC 14018 exhibited restriction activity, as the plasmid pKO403-*lacZ′-*Sp was digested into multiple fragments (Figure 1B). In contrast, the supernatant from *G. vaginalis* ATCC 49145 did not digest the plasmid DNA (Figure 1B). Based on these results, we selected *G. vaginalis* ATCC 49145 for transformation protocol development.

Since there are no reported plasmids in *Gardnerella*, we used the *Bifidobacterium-E. coli* shuttle vector, pKO403-*lacZ′-*Sp, which contains a spectinomycin resistance gene for the selection of transformants. To assess spectinomycin as a selection method, its MIC for *G. vaginalis* ATCC 41945 was determined in an agar dilution assay. *G. vaginalis* ATCC 49145 was susceptible to spectinomycin with an MIC of 2 µg/ml.

### Competent cell preparation and electroporation

Multiple protocols were evaluated for creation of electrocompetent *G. vaginalis* ATCC 49145. Bacteria for electrocompetent cell preparation were collected from log phase cultures grown in mBHI with 200 mM glycine. The inclusion of glycine was necessary to prevent clumping of bacteria during the washing steps (18). Parameters investigated included the wash solution, amount of plasmid DNA (40-600 ng), electroporation parameters (voltage, resistance, fresh or frozen competent cells) and recovery time (1.5 h or overnight (15-18 h)) (Supplemental Table 1). Results of each protocol were evaluated by plating electroporated bacteria on CBA plates containing spectinomycin ranging from 4-16 µg/ml for the selection of transformants carrying pKO403-*lacZ*′-Sp.

Bacteria were washed with water followed by two washes of 10 % glycerol, three washes with SHMG, or three washes with sucrose citrate buffer. Effects of wash solutions on viability was evaluated by plating bacteria on CBA before and after washing. The percentage of viable cells recovered after washing with water followed by 10 % glycerol, SHMG, or sucrose citrate buffer was 84 %, 94 % and 98 %, respectively, indicating that none of the washing solutions had a substantial impact on the viability of the bacteria. Therefore, we proceeded to electroporate competent cells made with all three wash solutions, assuming that they might yield different transformation efficiencies regardless of the initial effects on viability. Similarly, none of the voltages (1.5 kV, 1.8 kV, 2 kV or 2.5 kV) or resistance settings (200 Ω, 400 Ω, 800 Ω) were lethal and all the combinations tested resulted in time constants ranging from 4 ms to 18 ms (20). Spectinomycin-resistant transformants were not recovered using any of the tested protocols until we used fresh competent cells electroporated immediately following preparation and extended the recovery time to an overnight incubation (15-18 h). Use of frozen competent cells and recovery times of less than 6 hours did not result in successful transformation. Small numbers of transformants (<10 cfu) were recovered when competent cells were made by washing bacteria with water in the initial wash, followed by two washes with 10 % glycerol at either room temperature or on ice, and freshly made competent cells and 400 ng of plasmid DNA were electroporated at 1.5 kV, 800 Ω or 2.5 kV, 200 Ω with overnight recovery in mBHI. The efficiency of the transformation protocol developed was 2.5-4.5×10^2^ cfu/µg DNA indicating very low efficiency; however, it was reproducible. Addition of dimethyl sulfoxide (DMSO) has been reported to improve transformation efficiency (30), however, in this study, adding 5 % DMSO to the recovery medium did not enhance transformation efficiency.

### Recovery of plasmid pKO403-*lacZ′*-Sp from transformants

The next step was to confirm that the bacterial colonies that grew on the selective media after electroporation were due to the presence of plasmid pKO403-*lacZ′-*Sp. Direct colony PCR on nine colonies targeting the Sp^R^ gene resulted in the expected 843 bp amplicon. Four colonies were selected to inoculate broth cultures (mBHI with 16 µg/ml spectinomycin) and plasmid purification from cell pellets was attempted using the QIAprep Spin Miniprep Kit. Nanodrop quantification of the extracted plasmid from *Gardnerella* transformants showed concentrations of 45-305 ng/µl and an A_260_/A_280_ ratio of 1.9 indicating high quality DNA in all four extractions.

The nucleotide sequence of the plasmid isolated from *G. vaginalis* ATCC 49145 was 100 % identical sequence of pKO403-*lacZ′-*Sp provided by the supplier (Addgene Cat# 174726).

To confirm that low transformation efficiency was not the result of an undetected restriction activity, we performed additional electroporations using plasmid DNA extracted from *G. vaginalis* transformants instead of plasmid DNA extracted from *E. coli*. No improvement in transformation efficiency was observed.

### Spectinomycin susceptibility of transformant

To quantify the difference in spectinomycin susceptibility between wildtype *G. vaginalis* ATCC 49145 and the transformant, MICs were determined using agar dilution and broth microdilution assays. Growth of *G. vaginalis* ATCC 49145 with pKO403-*lacZ′-*Sp was observed up to 512 µg/ml in both agar and broth assays, whereas no growth of the wildtype strain was observed at any of the spectinomycin concentrations tested (Table 1). The growth patterns of the transformant and the wild-type strain were identical in both agar and broth cultures, although the transformant colonies at 512 μg/ml were small compared to the typical *Gardnerella* colonies.

**Table 1.** Spectinomycin tolerance of *G. vaginalis* ATCC 49145 and *G. vaginalis* ATCC 49145 with the pKO403-*lacZ′-*Sp plasmid in agar and broth.

|  | CBA plates |  |  |  |  |  |  | Modified BHI broth |  |  |  |  |  |  |
| --- | --- | --- | --- | --- | --- | --- | --- | --- | --- | --- | --- | --- | --- | --- |
| Sp (µg/ml) | 0 | 16 | 32 | 64 | 128 | 256 | 512 | 0 | 16 | 32 | 64 | 128 | 256 | 512 |
| Wildtype | +<sup>1</sup> | -<sup>2</sup> | - | - | - | - | - | + | - | - | - | - | - | - |
| Transformant | + | + | + | + | + | + | +<sup>3</sup> | + | + | + | + | + | + | + |

<sup>1</sup>Presence of growth, <sup>2</sup>Absence of growth, <sup>3</sup>Tiny colonies

### Plasmid copy number

Plasmid copy number for a particular plasmid can vary depending on the host species (31). To determine the copy number for pKO403-*lacZ′-*Sp in *G. vaginalis* ATCC 49145, quantitative PCR was done using a calibrator plasmid as described in the methods. Using the calibrator plasmid as a template, the optimal annealing temperature for the PCR assays targeting both the chromosomal uppS gene and the plasmid Sp^R^ gene was 57.5 °C and efficiencies of the PCR assays were 95.4 % and 90.0 %, respectively (Supplemental Figure 1). Real-time quantitative PCR was performed on total DNA (1:10 and 1:100 dilutions) from a *G. vaginalis* transformant using both PCR assays and Cq values were used to calculate copy numbers of uppS (chromosome) and Sp^R^ gene (plasmid) per reaction based on the calibrator standard curves. Results from the two template dilutions were consistent, and the plasmid copy number per chromosome copy was determined to be 2.72 (average of two experiments) (Table 2).

**Table 1.**
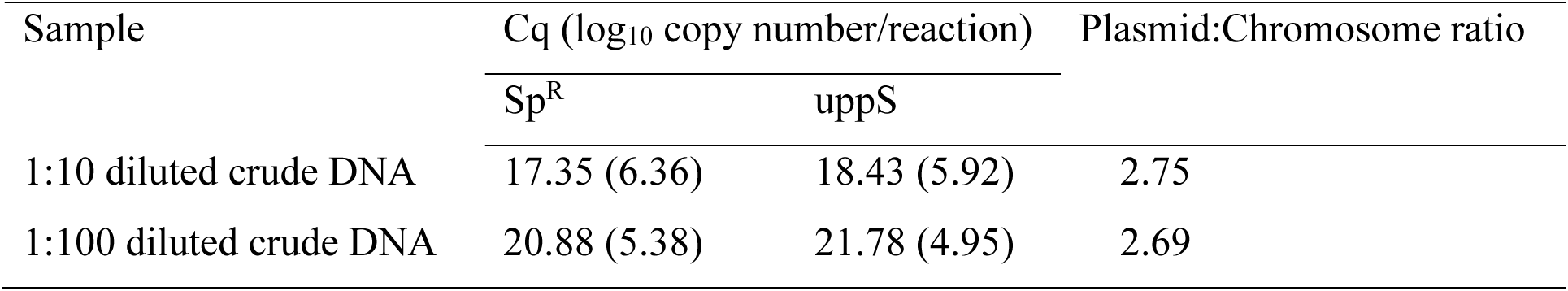
Plasmid copy number calculation.

| Sample | Cq (log<sub>10</sub> copy number/reaction) |  | Plasmid:Chromosome ratio |
| --- | --- | --- | --- |
|  | *Sp<sup>R</sup>* | *uppS* |  |
| 1:10 diluted crude DNA | 17.35 (6.36) | 18.43 (5.92) | 2.75 |
| 1:100 diluted crude DNA | 20.88 (5.38) | 21.78 (4.95) | 2.69 |

### β-galactosidase activity of *Gardnerella* transformants

*G. vaginalis* is β-galactosidase positive by definition (32), but if the β-galactosidase gene on pKO403-*lacZ′-*Sp was being expressed, the transformants would have greater enzymatic activity than the wildtype. When the *G. vaginalis* ATCC 49145 wildtype and the *G. vaginalis* ATCC 49145 with pKO403-*lacZ′*-Sp were grown in the presence of X-gal, blue colonies were observed in both the strains; however, the intensity of the colour of the transformants was greater than that of the wild type, consistent with a higher level of β-galactosidase activity in the transformed cells (Figure 2).

**Figure 2.**
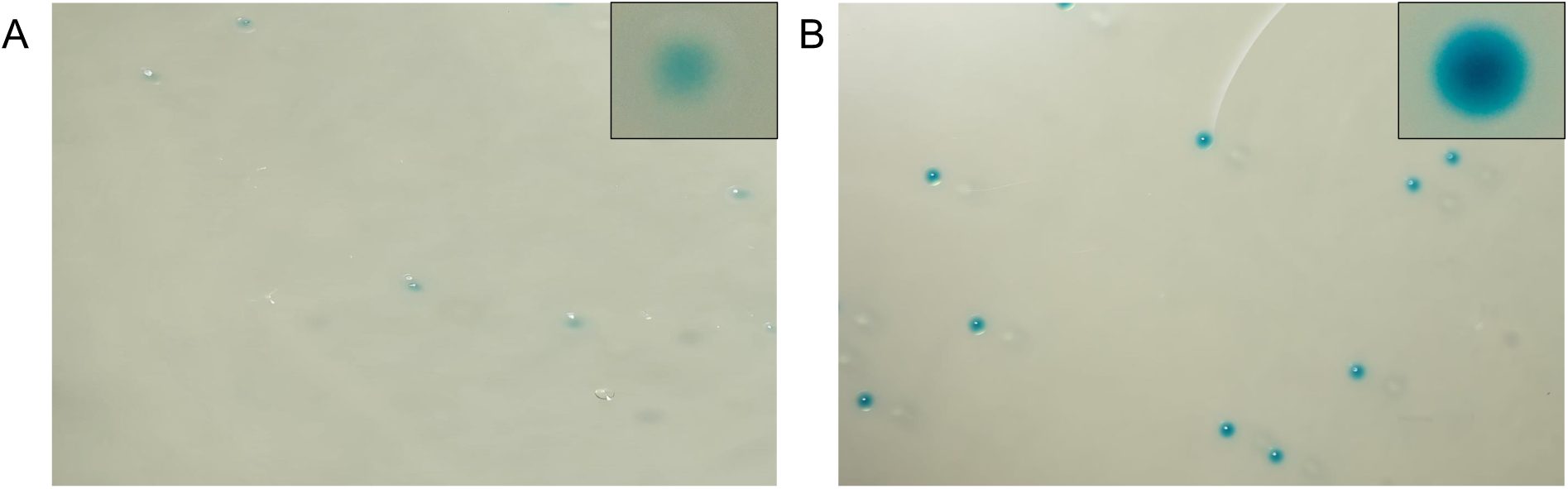
A) *G. vaginalis* grown in the presence of X-gal on MHA supplemented with horse serum. B) *G. vaginalis* + pKO403-*lacZ′-*Sp in the presence of X-gal on MHA supplemented with horse serum and 50 μg/ml of spectinomycin. The inset in the top right corner of each panel shows an enlarged single colony. Image taken from Canon R6 with Laowa 100mm f/2.8 2× Ultra Macro Apo lens.

### Plasmid stability

Plasmid stability refers to how effectively daughter cells inherit the plasmid during cell replication (33). We evaluated the stability of pKO403-*lacZ′-*Sp over 18 h to 24 h at which point they reach stationary phase (Figure 1). When *G. vaginalis* ATCC 49145 with the plasmid was grown with spectinomycin, 53.0 % of colony forming units were spectinomycin resistant at 18 h and 63.3 % at 24 h (Figure 3). In the absence of selection, the proportion of spectinomycin resistant colony forming units was only 12.4 % at 18 h and further reduced to 5.0 % by 24 h (Figure 3).

**Figure 3.**
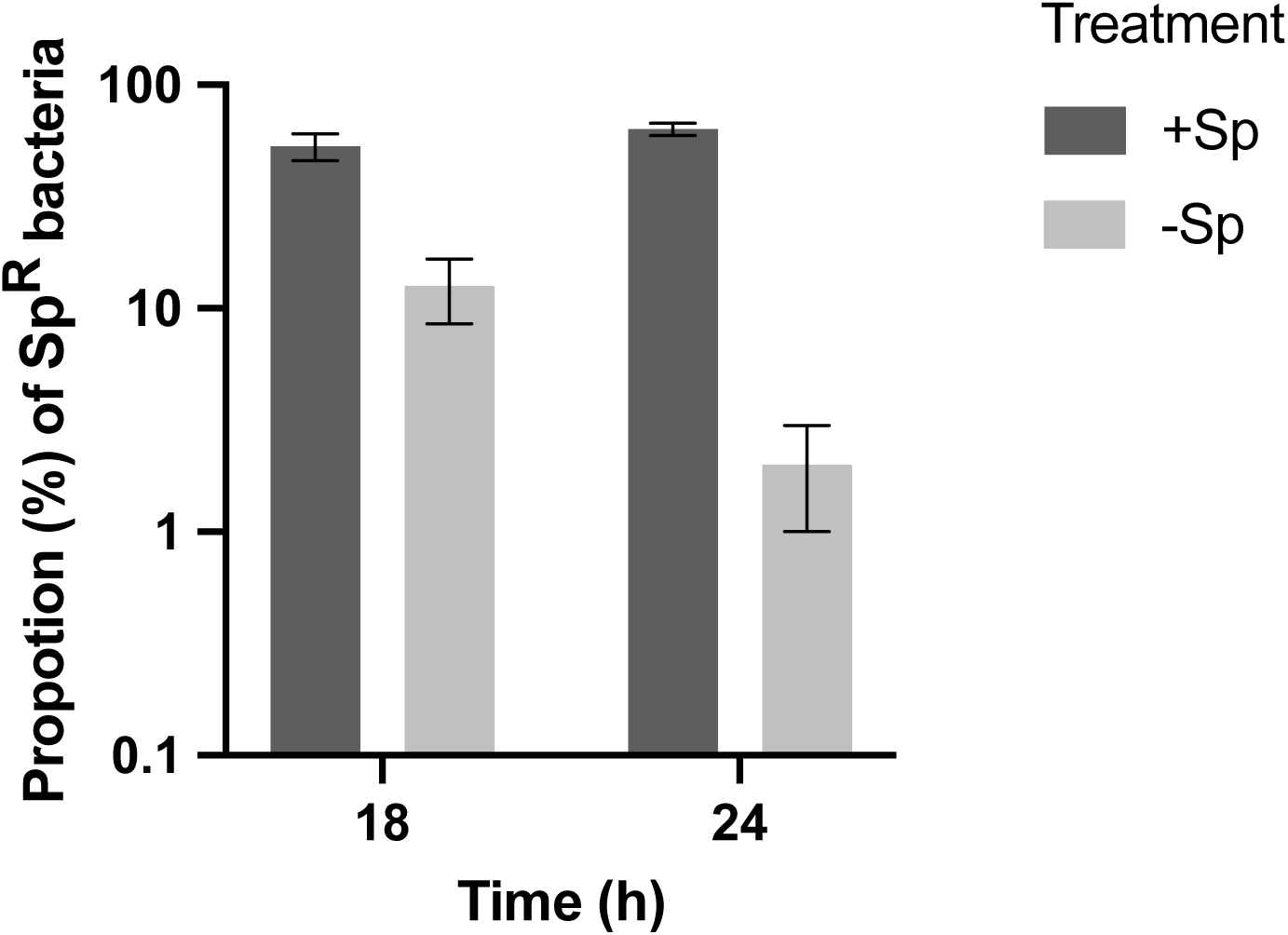
Plasmid stability: *G. vaginalis* with pKO403-*lacZ′-*Sp was grown in the presence or absence of spectinomycin (as indicated in the legend) for 18 h or 24 h and then plated in CBA with and without spectinomycin. Colony-forming units on selective and non-selective media were counted to calculate the proportion of Sp^R^ bacteria. Each bar represents the average of three replicates, and error bars indicate the standard deviation.

Plasmid pKO403-*lacZ′-*Sp is known to be cured at 42 °C in *Bifidobacterium* due to a temperature sensitive mutation in the pTB6 original of replication (27). Prior to evaluating the curing process in *G. vaginalis*, we first confirmed the ability of wildtype *G. vaginalis* ATCC 49145 to grow at the non-permissive temperature on CBA and in mBHI broth. The next step was to evaluate the growth of *G. vaginalis* ATCC 49145 with the plasmid at 42 °C. We found that the transformants did not grow in the mBHI broth at 42 °C regardless of the presence or absence of selection, which negated any further experiments on plasmid curing. We did, however, observe atypical, tiny colonies of transformants at 42 °C on CBA both with and without selection.

## Discussion

*Gardnerella* species play a major role in the pathogenesis of bacterial vaginosis (34–37). Although an abundance of *Gardnerella* is considered a primary initiator of this condition, the absence of genetic tools has hindered mechanistic research necessary to explore its biology, physiology, and ecological role in bacterial vaginosis. Since this organism was first described by Leopold in 1953 (38) and named by Gardner and Dukes in 1955 (39) to our knowledge, no genetic tools have been developed for *Gardnerella*. Thus, this study aimed to establish a protocol for transforming *Gardnerella vaginalis* through electroporation and to characterize transformants carrying a *Bifidobacterium-E. coli* shuttle vector.

Our selection of a candidate strain of *G. vaginalis* was based on identifying an isolate that exhibited a relatively high growth rate without requiring anaerobic incubation. Not surprisingly, we found that the *G. vaginalis* type strain ATCC 14018 and *G. vaginalis* ATCC 49145 grew more than clinical isolates of other species in the provided growth conditions (Figure 1A). The ATCC strains of *G. vaginalis* are known to grow well in 7% CO_2_ and are highly laboratory-adapted (40). Abundant growth in simple conditions is an advantage when large numbers of bacteria are needed. However, adaptation of transformation protocols for non-laboratory adapted strains and other species in the genus will be a focus for future work. Ideally, the candidate strains should also lack restriction activity; a major barrier to introducing foreign DNA (12). The observed restriction activity in this study for *G. vaginalis* ATCC 14018 and corresponding lack of restriction activity in *G. vaginalis* ATCC 49145 was concordant with previously published observations (Figure 1B) (41, 42). Strain level variation in restriction modification systems has been reported in other species including *Helicobacter pylori* and *Neisseria gonorrhoeae* (43, 44), which emphasizes the importance of evaluating individual strains.

As *G. vaginalis* ATCC 49145 is a non-model organism, transformation is a challenging process. *G. vaginalis* has a Gram-positive cell wall although its very thin peptidoglycan layer results in its description as “Gram variable” (45). Methods for inducing competence in Gram-positive bacteria including *Bifidobacterium*, *Staphylococcus*, *Listeria* and *Bacillus* are widely reported (12, 46–48) and so this type of cell wall does not present an insurmountable barrier to transformation. We harvested bacteria for competent cell preparation in mid exponential phase as is done in most transformation protocols. Interestingly, bacteria that are harvested from earlier growth stages (49) or during stationary phase (50) have been reported to yield higher efficiency competent cells and so this aspect of inducing competence in *G. vaginalis* should be explored in future studies.

During *G. vaginalis* ATCC 49145 competent cell preparation, we experimented with 10% glycerol (12, 47, 51), SHMG (12, 46) and sucrose citrate buffer (52) as wash solutions since they have been used successfully in making other bacteria competent. Regardless of the wash solution, a major issue encountered with *G. vaginalis* was aggregation. To overcome this, glycine was added to the broth culture (18). Glycine contributes to weakening the cell wall by interfering with the peptidoglycan synthesis (53) and is known to increase transformation efficiency by facilitating foreign DNA introduction (54). None of the wash solutions we evaluated had a significantly negative impact on viability and could be considered for use in making competent *G. vaginalis*. Wash temperature has been reported to affect competent cell preparation and transformation efficiency (55, 56), however we found no difference in the transformation efficiency of *G. vaginalis* ATCC 49145 washed either in ice cold or room temperature wash solutions.

Plasmids are naturally present in many species across the prokaryotic tree of life, offering advantages in ecology and evolution by facilitating the transfer of beneficial traits necessary for survival in complex communities and harsh environments (57). Some bacteria, including *Brucella abortus* and *Bartonella bacilliformis*, have not exhibited any naturally occurring plasmids, which may be due to their associated fitness cost (58). However, *B. abortus* has been successfully transformed and can retain broad host range plasmid pTH10, reinforcing the idea that bacteria like *G. vaginalis* that lack natural plasmids can effectively maintain introduced plasmids. We chose *Bifidobacterium - E. coli* shuttle vector, pKO403-*lacZ′*-Sp (27) for our experiments since *Bifidobacterium* and *Gardnerella* are taxonomically related (family Bifidobacteriaceae) and we speculated that the pTB6 origin of replication would support replication in *G. vaginalis*. Selection based on spectinomycin resistance was also feasible given the low MIC for this antibiotic we measured for *G. vaginalis* (2 µg/ml).

The voltage, capacitance, and resistance during electroporation influence both the effective delivery of nucleic acid to cells and cell viability. Voltages ranging from 1.5 kV to 2.5 kV have been widely used in bacterial transformation (14, 16, 49, 59, 60). Transformation efficiency may also be affected by pulse length (59). We did not observe any differences in transformation efficiency of *G. vaginalis* with the different voltages and pulse lengths that were investigated in this study (Supplemental Table 1). Successful transformations were achieved with large amounts of plasmid DNA (up to 600 ng) in the electroporation, similar to what has been reported for *Legionella pneumophila* where 200 ng of plasmid was routinely included in electroporations (59). Further experiments would be needed to determine the relationship between amount of DNA and transformation efficiency.

Following electroporation, a recovery phase in rich medium is required to allow expression of antibiotic resistance genes prior to plating on selective media. While recovery times of less than an hour may be sufficient for rapidly growing model organisms such as *E. coli*, slow growing bacteria like *Mycobacterium tuberculosis* require a prolonged period of recovery (up to 16 h) to enhance transformation efficiency (61). We found that recovery times at least 15 h were essential in obtaining transformants of *G. vaginalis* ATCC 49145. The addition of DMSO to the recovery medium increases the transformation efficiency of bacteria such as *Agrobacterium* since it acts as a permeability enhancer and promotes membrane fusion (30). Addition of DMSO to the mBHI recovery medium in our experiments had no apparent benefit.

The protocol that was developed in this study resulted in the recovery of transformants with an efficiency of ∼10² cfu/µg of DNA, which is very low but was reproducible. To enhance efficiency, various alternative cell-weakening agents, including DL-threonine, lysozyme, and penicillin G (62, 63) may be explored. Additionally, it would be beneficial to collect cells at different stages of the growth cycle and evaluate the transformation efficiency. Since plasmid size and type may play an important role in determining transformation efficiencies (64), other plasmids from bifidobacteria are worth exploring. We did rule out the possibility that an undetected restriction activity was responsible for the lower transformation efficiency since the results of electroporation with plasmid DNA extracted from *G. vaginalis* ATCC 49145 transformants were no more efficient than that observed when transforming with pKO403-*lacZ′-* Sp extracted from *E. coli*.

In the resulting transformants, the plasmid copy number was calculated in *G. vaginalis* ATCC 49145 relative to the chromosome number since the ploidy of bacteria varies based on strain and growth stage (65). The pKO403-*lacZ′-*Sp copy number was ∼3, which is considered low copy number (33). Nevertheless, the gene dosage was sufficient to increase the MIC of spectinomycin to at least 512 µg/ml (Table 1) and substantially increase β-galactosidase activity (Figure 2); observations that also confirm the suitability of promoters in pKO403-*lacZ′-*Sp for gene expression in *G. vaginalis*.

In the absence of selection, plasmids impose a fitness cost on their hosts and so may be rapidly lost if the genes they carry are not needed for survival (66). In the absence of spectinomycin selection, *G. vaginalis* transformants rapidly lost the pKO403-*lacZ′-*Sp plasmid with only 5% of bacteria retaining their Sp^R^ phenotype after 24 hours (Figure 3). The proportion of Sp^R^ bacteria recovered after 24 hours of growth with selection was markedly higher (63%) but this result is consistent with the plasmid being inherently unstable in *G. vaginalis*. Instability even in the presence of selection has also been reported for derivatives of pKO403 in *Bifidobacterium longum* (27). For *G. vaginalis* this observation is likely related to whatever factors contribute to an absence of naturally occurring plasmids in this species. If plasmid stability is high, curing of plasmids in some genetic engineering approaches may be challenging. For this reason, the pKO403-*lacZ′-*Sp plasmid was designed to provide a temperature-sensitive origin of replication to facilitate curing. We were unable to investigate this feature since transformants were not viable under selection at the non-permissive temperature, likely due to the combined fitness cost of the plasmid and the temperature stress.

Overall, our findings demonstrate the feasibility of low-efficiency transformation of *G. vaginalis* ATCC 49145 by electroporation and that plasmids compatible with bifidobacteria such as pKO403-*lacZ′-*Sp offer a starting point for the development of genetic tools for the manipulation of *G. vaginalis* and potentially other species of this clinically significant genus.

## Materials and methods

### Strains and plasmid

*G. vaginalis* ATCC 49145 and *G. vaginalis* ATCC 14018 were purchased from the American Type Culture Collection (ATCC). *Gardnerella swidsinskii* (LP 23b), *Gardnerella piotii* (LP 13C) and *Gardnerella leopoldii* (LP 57) were isolated from previously described vaginal swab samples (67). Plasmid pKO403-*lacZ′-*Sp was purchased from Addgene (Cat# 174726) (27).

### Media and culture conditions

*Gardnerella* isolates were revived from −80 °C stocks on Columbia blood agar plates (BD BBL™, Fisher Scientific) and incubated at 37 °C, in 7 % CO_2_ for 48 h. For broth culture, *Gardnerella* isolates were grown in modified brain heart infusion broth (mBHI), which is BHI supplemented with 0.25% (w/v) maltose and 10% (v/v) heat-inactivated horse serum (Gibco). *G. vaginalis* ATCC 49145 with plasmid pKO403-*lacZ*′-Sp was grown in the presence of 50 µg/ml of spectinomycin in CBA and mBHI unless otherwise indicated. To assess β-galactosidase activity, isolates were grown on Mueller Hinton agar, containing 10% (v/v) heat-inactivated horse serum and X-gal (20 mg/ml), with or without spectinomycin.

To quantify bacteria (cfu/ml), track-dilution was performed from dilution series prepared from broth cultures, and 8 µl of each dilution was spotted onto CBA or CBA with 50 µg/ml spectinomycin depending on the experiment (68).

### Determination of MIC of spectinomycin

The minimum inhibitory concentration (MIC) of spectinomycin for *G. vaginalis* ATCC 49145 was determined by the agar dilution method. A suspension of bacteria from CBA plates (0.5 McFarland standard) was prepared in 5 ml of distilled water. Inoculum spots of 2 µl were then placed on prepared antibiotic plates (Mueller Hinton agar with 0.125 µg/ml to 128 µg/ml spectinomycin), and the plates were incubated at 37 °C, in 7 % CO_2_ for 48 h. The MIC was determined as the lowest concentration at which bacterial growth was not observed.

To determine the spectinomycin tolerance level of *G. vaginalis* ATCC 49145 transformed with pKO403-*lacZ*′-Sp relative to wildtype *G. vaginalis* ATCC 49145, bacteria were grown in triplicate on agar (CBA with 0-512 µg/ml spectinomycin) and in broth (mBHI with 0-512 µg/ml spectinomycin). Both agar and broth were incubated for 48 h at 37 °C in 7 % CO_2_.

### Assessment of restriction activity

Isolates were grown in mBHI until they reached OD_600_ ∼0.9. Subsequently, 1 ml of this broth was pelleted by centrifugation at 16800 × *g* for 2 min and the supernatant was filtered through a 0.2 µm Acrodisc syringe filter (Cytiva Life Sciences). Afterwards, 2 µl of the filtered supernatant was mixed with 8 µl of pKO403-*lacZ*′-Sp (135 ng/µl) in the presence of 0.1 µg/µl RNase A (ThermoFisher Scientific), and 2 µl of CutSmart buffer (New England Biolabs). The reaction mixture was incubated for one hour at 37 °C, and the resulting products were analyzed by electrophoresis using a 1% agarose gel.

### Preparation of competent cells and electroporation

*G. vaginalis* ATCC 49145 was revived from −80 °C on CBA plates at 37 °C for 24 h in 7 % CO_2_. A 10 µl inoculation loop was used to collect colonies and transfer to 5 ml of mBHI for incubation at 37 °C for 12 h in 7 % CO_2_. The resulting broth culture was then diluted with 55 ml of mBHI medium containing 200 mM glycine and incubated at 37 °C in 7 % CO_2_. After the culture reached an OD_600_ of 0.6-0.8, it was placed in an ice bath for 30 minutes. Next, 50 ml of the broth culture was transferred to an ice-cold tube and centrifuged at 1000 × *g* for 15 min at 4 °C. The supernatant was decanted, and the remaining cell pellet was washed with 25 ml of ice-cold wash solution by gently pipetting up and down, followed by centrifugation at 1000 × *g* for 20 min. Wash solutions evaluated during protocol development were 10 % glycerol, SHMG (0.25 M Sucrose, 1 mM HEPES, 1 mM MgCl_2_, 10 % Glycerol), sucrose citrate buffer (0.5 M sucrose, 1 mM citrate), and water. Following the first wash, the supernatant was decanted, and the cells were resuspended in 12.5 ml of wash solution and centrifuged at 1000 × *g* for 20 min. The resulting pellet was resuspended in 0.5 ml of ice-cold wash solutions and centrifuged at 1000 × *g* for 20 min. The final pellet was resuspended in 120 µl of 10 % glycerol and either frozen at −80 °C or used immediately.

Freshly prepared or frozen competent cells (40 µl) were mixed with 2 µl of plasmid pKO403-*lacZ*′-Sp (40-600 ng) in a microcentrifuge tube and kept on ice for 1 minute. The mixture was then transferred to a 0.2 cm cuvette (Bio-Rad) and electroporated using a Bio-Rad Gene Pulser at 1.5-2.5 kV, 200-800 Ω and 25 µF. Immediately after electroporation, 1 ml of mBHI was added to the cuvette as the recovery medium. The entire suspension was transferred to a 10 ml sterile culture tube and incubated for 1 to 4 h or overnight at 37 °C in 7 % CO_2_. Following incubation, 100 µl of the culture was spread on each of eight CBA plates with spectinomycin ranging from 2-50 µg/ml, and the remaining 200 µl of the inoculum was spread on CBA plates without spectinomycin to check the viability of the bacteria. Plates were incubated for 48-72 h at 37 °C in 7 % CO_2_.

### Polymerase chain reaction for detection of plasmids

To confirm the presence of the plasmid in electroporated *G. vaginalis*, colony PCR was performed targeting the spectinomycin resistance gene of pKO403-*lacZ*′-Sp. Primers were designed using Primer3Plus (version:3.3.0). Each PCR reaction was 25 µl in total volume and contained 0.4 µM of each of forward primer 5’-TCA AAA TAG TGA GGA GGA TAT ATT TGA-3’ and reverse primer 5’-CTG AAT TCC CAT TAA ATA ATA AAA CAA-3’, 1× PCR buffer, 2.5 mM MgCl_2_, 0.2 mM dNTPs, 0.05 U Taq DNA Polymerase. Thermocycling was carried out at 94 °C for 5 min followed by 40 cycles of denaturing at 94 °C for 30 sec, annealing at 55 °C for 40 sec, and extension at 72 °C for 45 sec with final extension for 10 min at 72 °C. The resulting PCR product were analyzed by electrophoresis on a 0.8% agarose gel.

### Plasmid extraction purification and sequencing

Putative transformants were grown in mBHI in the presence of 16 µg/ml of spectinomycin and plasmid extraction was performed using a commercial kit (QiAprep Spin Miniprep Kit). The concentration and purity (A_260_/A_280_ ratio) of the purified plasmid DNA was evaluated by spectrophotometry (Nanodrop 2000c) and the purified plasmid was sequenced using Oxford Nanopore Technology (Plasmidsaurus, San Francisco, CA).

### Plasmid copy number calculation

To estimate the plasmid copy number of pKO403-*lacZ*′-Sp in transformants, the ratio of plasmid copy number to chromosome copy number was determined (69). Quantitative real-time PCR was performed with primers targeting the uppS (polyprenyl diphosphate synthase) gene (JH0931: 5’-TTA GTG ATC CTA GCC GCG TG-3’ and JH0932: 5’-GCC GTC CAT AAT CAC GCC TA-3’) on the *G. vaginalis* ATCC 49145 chromosome and the Sp^R^ gene (JH0949: 5’-TCA GGA TGA TGA AAC CAA CTC T-3’ and JH0950: 5’-ACG AAC TGC TAA CAA AAT TCT CTC C-3’) of pKO403-*lacZ′-*Sp. To determine the optimal annealing temperature for the PCR assays, conventional PCR with an annealing temperature gradient was performed. A standard curve was constructed for plasmid copy number determination using a 10-fold dilution series of a calibrator plasmid (10^7^ to 10^2^ copies per reaction). The calibrator is a synthetic construct that incorporates the products of both the uppS primers and the Sp^R^ gene primers, along with their binding sites, ligated into the pIDTSmart ampicillin vector (synthesized by IDT) (Supplemental Figure 2).

Template DNA for the qPCR was prepared by suspending three to four loopful (10 µl) of *G. vaginalis* ATCC 49145+pKO403-*lacZ*′-Sp in 250 µl molecular biology grade water followed by incubation at 99 °C for 3 min in a heat block and pelleting of cellular debris by centrifugation at 4000 × g for 4 min. The clarified supernatant was diluted 1:10 or 1:100 and 2 μl was used as a template in real-time quantitative PCR. The master mix of 10 μl was prepared for qPCR with 5 μl of 2× SYBR mix (Biorad-IQ TM SYBR green supermix), 0.4 μl of each forward and the reverse primer (10 pmol/μl) and 2.2 μl of molecular biology grade water. PCR was performed using the Bio-rad CFX connect real time system. The cycling parameters were 95 °C 3 min followed by 40 cycles of 10 s at 95 °C, 10 s at 57.5 °C and 15 s at 72 °C. Each PCR reaction was performed in duplicate. *G. vaginalis* ATCC 49145 wild type was used as a positive control for uppS gene and the plasmid pKO403-*lacZ*′-Sp was used as the positive control for the Sp^R^ gene. No template controls were included with each run.

### Plasmid stability

*G. vaginalis* ATCC 49145 with the plasmid was revived from −80 °C on CBA with 50 µg/ml spectinomycin at 37 °C, in 7 % CO_2_ for 24 h. Two to three loopful of bacterial colonies from 10 µl inoculating loop were transferred to 3 ml of mBHI with 50 µg/ml spectinomycin and incubated at 37 °C, in 7 % CO_2_ until it reached OD600 ∼0.65. Next, 600 µl of the above broth was added to 30 ml mBHI with or without spectinomycin (50 µg/ml) and each suspension was divided into six polypropylene culture tubes such that each tube contained 5 ml. All 12 tubes were incubated at 37 °C, in 7 % CO_2_. After 18 h and 24 h of incubation, three replicate cultures were collected for each treatment (with or without spectinomycin) and track dilution was performed on CBA plates with or without 50 µg/ml spectinomycin and incubated at 37 °C, in 7 % CO_2_ for 48 h. The resulting bacterial colonies were counted to calculate the proportion of Sp^R^ bacteria (i.e. proportion of each culture carrying the plasmid) as follows: (CFU_CBA+spectinomycin_ / CFU_CBA_) × 100.

## Acknowledgements

This research was funded by a Natural Sciences and Engineering Research Council of Canada Discovery Grant to JEH. BMDNK received support from the Department of Veterinary Microbiology Student Support Fund. The authors are grateful to Champika Fernando for excellent technical support, and to Kosala Rajapaksa for assistance with photography.

---

**Supplemental Table 1:**
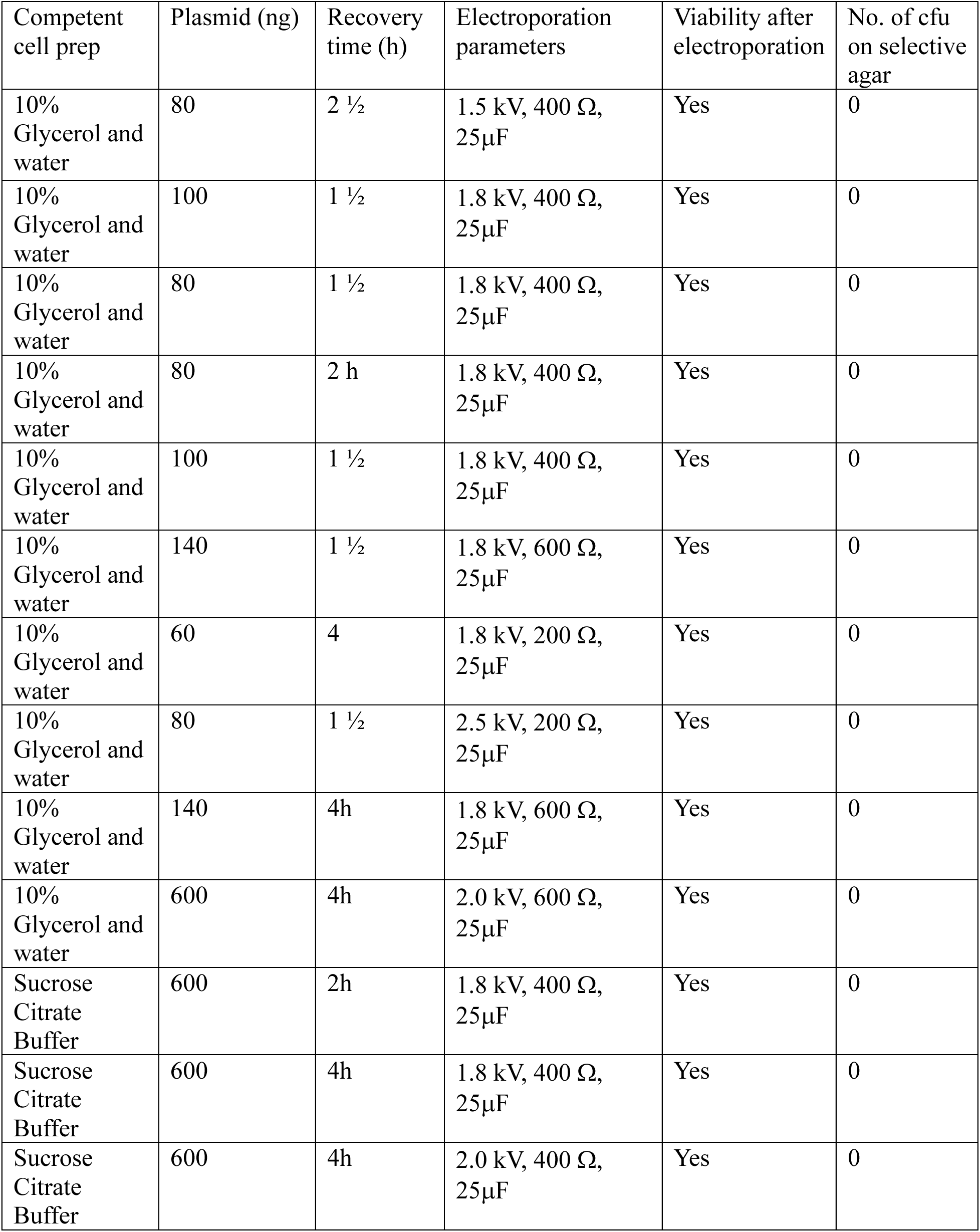
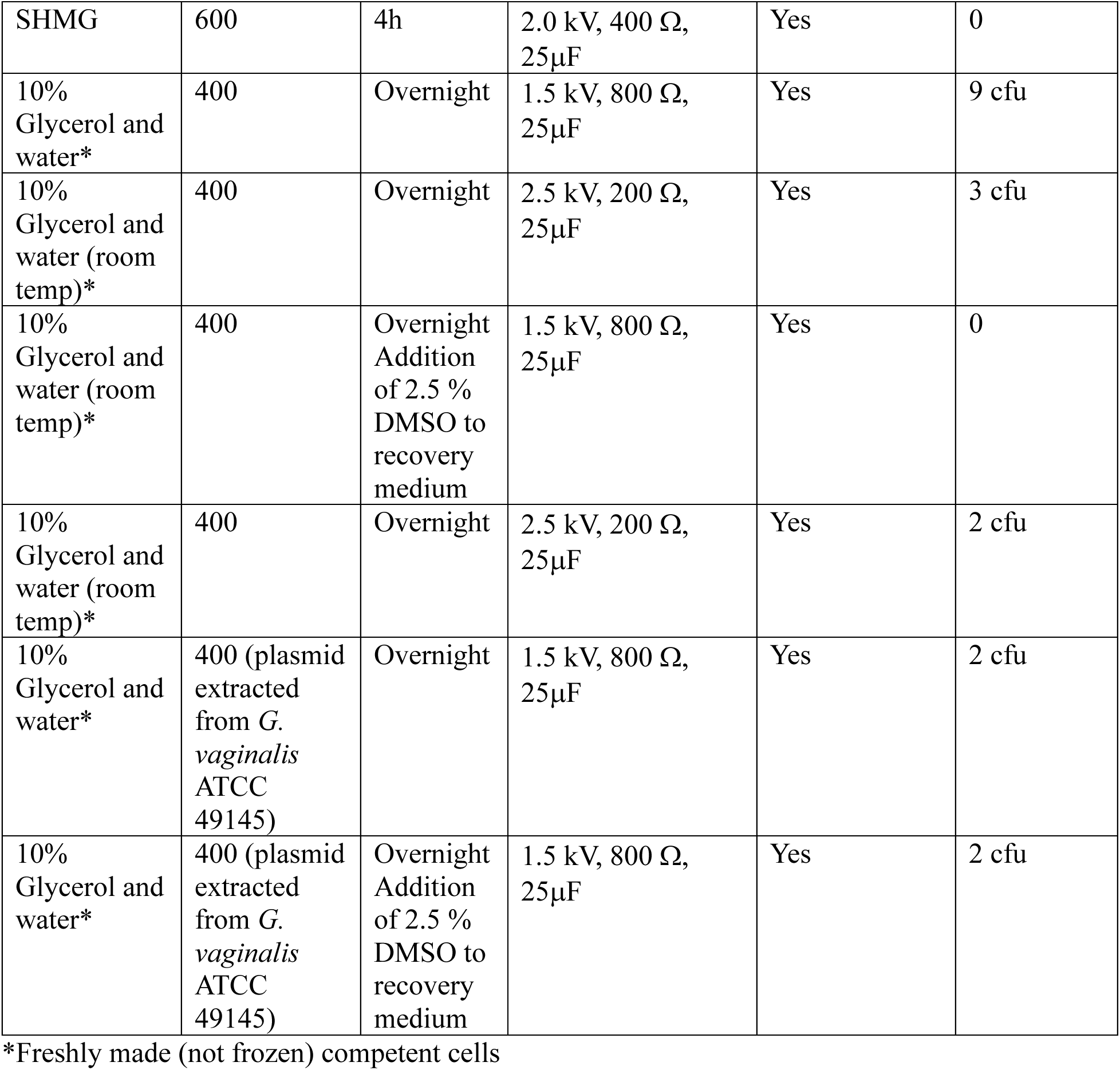
Results of transformation experiments in which various combinations of competent cell preparation, plasmid amount, electroporation parameters and recovery time were investigated.

| Competent cell prep | Plasmid (ng) | Recovery time (h) | Electroporation parameters | Viability after electroporation | No. of cfu on selective agar |
| --- | --- | --- | --- | --- | --- |
| 10% Glycerol and water | 80 | 2 ½ | 1.5 kV, 400 Ω, 25μF | Yes | 0 |
| 10% Glycerol and water | 100 | 1 ½ | 1.8 kV, 400 Ω, 25μF | Yes | 0 |
| 10% Glycerol and water | 80 | 1 ½ | 1.8 kV, 400 Ω, 25μF | Yes | 0 |
| 10% Glycerol and water | 80 | 2 h | 1.8 kV, 400 Ω, 25μF | Yes | 0 |
| 10% Glycerol and water | 100 | 1 ½ | 1.8 kV, 400 Ω, 25μF | Yes | 0 |
| 10% Glycerol and water | 140 | 1 ½ | 1.8 kV, 600 Ω, 25μF | Yes | 0 |
| 10% Glycerol and water | 60 | 4 | 1.8 kV, 200 Ω, 25μF | Yes | 0 |
| 10% Glycerol and water | 80 | 1 ½ | 2.5 kV, 200 Ω, 25μF | Yes | 0 |
| 10% Glycerol and water | 140 | 4h | 1.8 kV, 600 Ω, 25μF | Yes | 0 |
| 10% Glycerol and water | 600 | 4h | 2.0 kV, 600 Ω, 25μF | Yes | 0 |
| Sucrose Citrate Buffer | 600 | 2h | 1.8 kV, 400 Ω, 25μF | Yes | 0 |
| Sucrose Citrate Buffer | 600 | 4h | 1.8 kV, 400 Ω, 25μF | Yes | 0 |
| Sucrose Citrate Buffer | 600 | 4h | 2.0 kV, 400 Ω, 25μF | Yes | 0 |
| SHMG | 600 | 4h | 2.0 kV, 400 $\Omega$,<br>25μF | Yes | 0 |
| 10% Glycerol and water* | 400 | Overnight | 1.5 kV, 800 $\Omega$,<br>25μF | Yes | 9 cfu |
| 10% Glycerol and water (room temp)* | 400 | Overnight | 2.5 kV, 200 $\Omega$,<br>25μF | Yes | 3 cfu |
| 10% Glycerol and water (room temp)* | 400 | Overnight<br>Addition<br>of 2.5 %<br>DMSO to<br>recovery<br>medium | 1.5 kV, 800 $\Omega$,<br>25μF | Yes | 0 |
| 10% Glycerol and water (room temp)* | 400 | Overnight | 2.5 kV, 200 $\Omega$,<br>25μF | Yes | 2 cfu |
| 10% Glycerol and water* | 400 (plasmid<br>extracted<br>from *G.*<br>*vaginalis*<br>ATCC<br>49145) | Overnight | 1.5 kV, 800 $\Omega$,<br>25μF | Yes | 2 cfu |
| 10% Glycerol and water* | 400 (plasmid<br>extracted<br>from *G.*<br>*vaginalis*<br>ATCC<br>49145) | Overnight<br>Addition<br>of 2.5 %<br>DMSO to<br>recovery<br>medium | 1.5 kV, 800 $\Omega$,<br>25μF | Yes | 2 cfu |

\*Freshly made (not frozen) competent cells

**Supplemental Figure 1:**
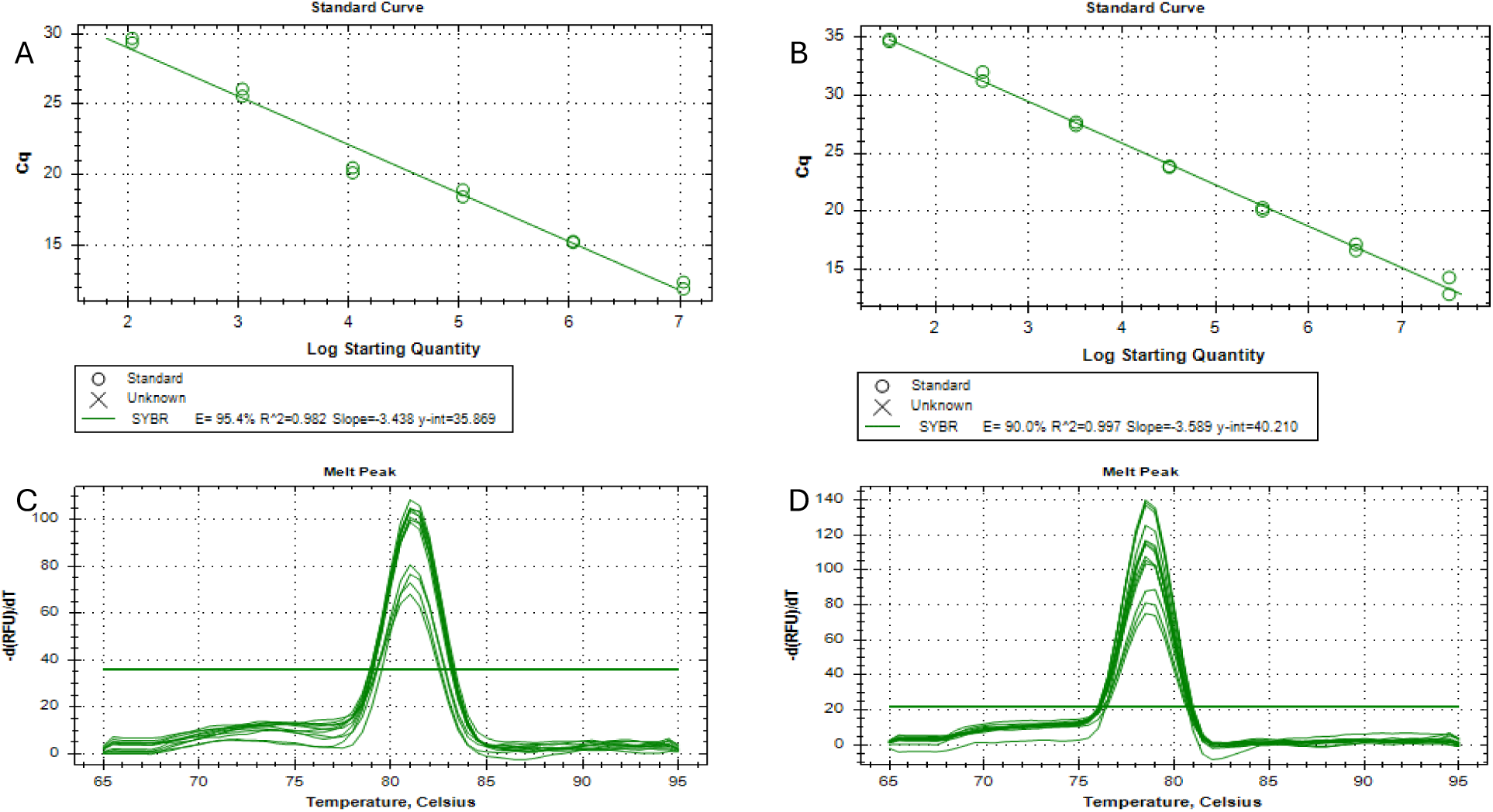
Performance of uppS and Sp^R^ gene SYBR green PCR assays. All reactions performed in duplicate using the calibrator plasmid as template. A) Standard curve for uppS assay, B) Standard curve for Sp^R^ gene assay, C) melt peak of the uppS PCR product, D) melt peak of the Sp^R^ gene product.

## Calibrator design

The following sequence shows the uppS gene fragment and the primer binding sites (JH0931 and JH0932 – highlighted in yellow) and the Sp^R^ gene fragment with primer binding sites (JH0949 and JH0950-highlighted in green). This 225 bp fragment was ligated into pIDTSmart Ampicillin (Supplemental Figure 2).

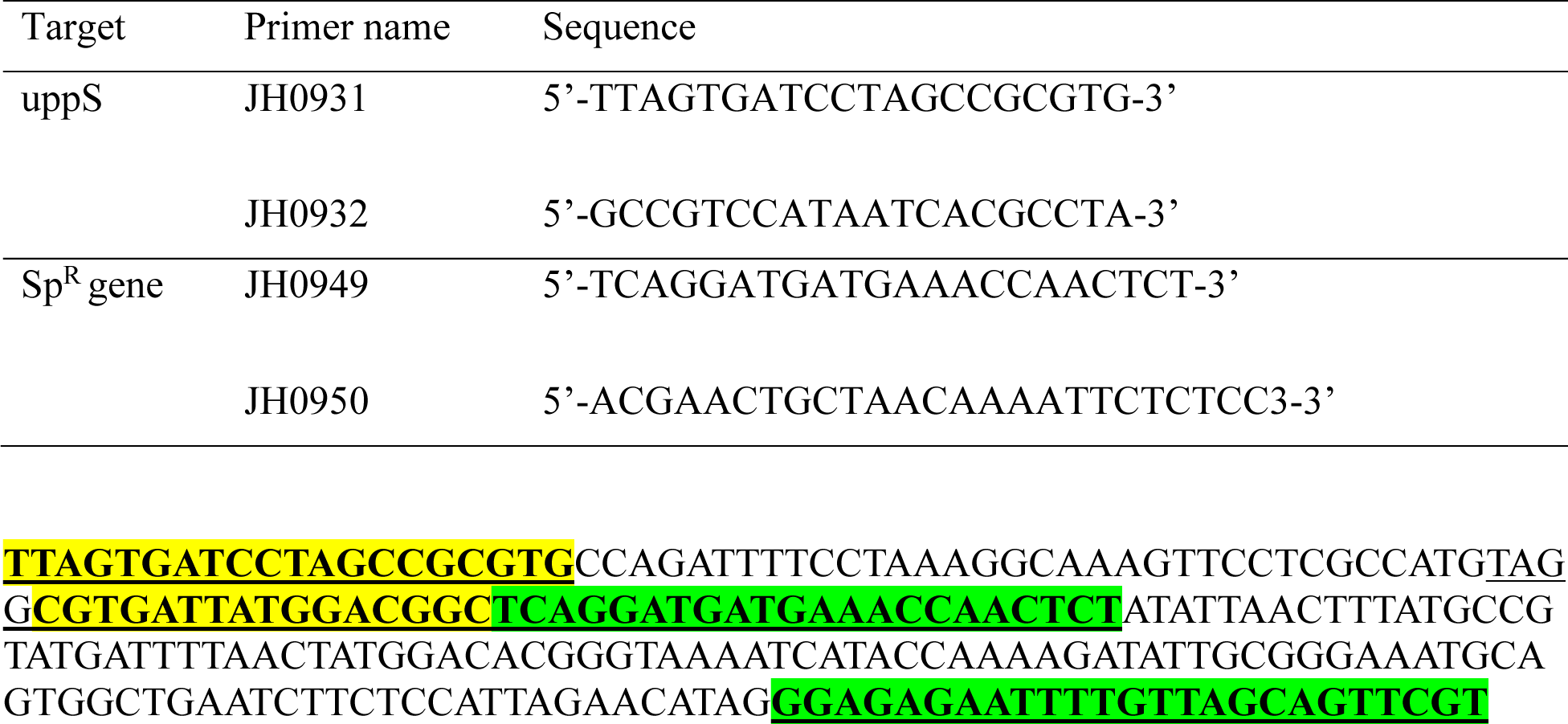

**Supplemental Figure 2:**
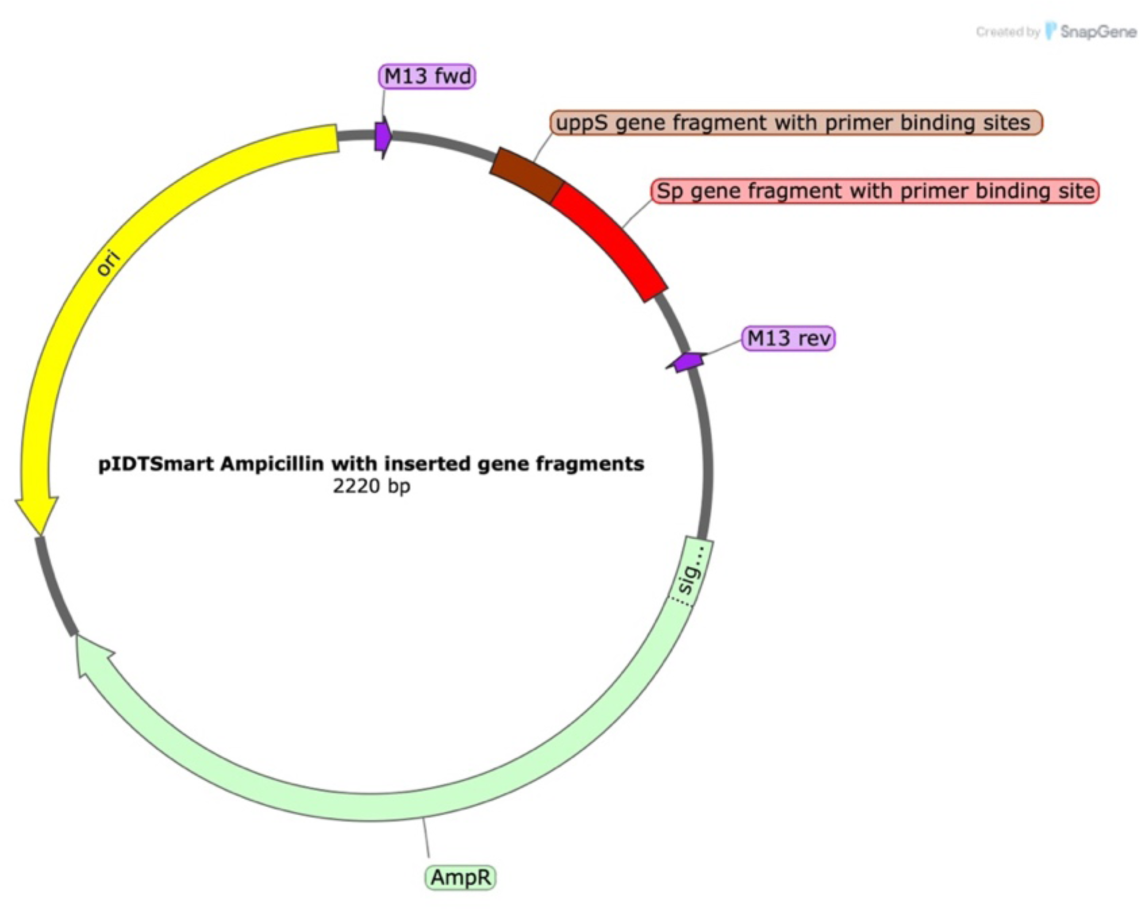
pIDTSmart Ampicillin plasmid with inserted uppS gene fragment and primer binding sites and Sp^R^ gene fragment with primer binding sites.

